# The role of theta and gamma oscillations in item memory, source memory, and memory confidence

**DOI:** 10.1101/2023.10.18.562880

**Authors:** Syanah C. Wynn, Christopher D. Townsend, Erika Nyhus

## Abstract

Theta and gamma oscillations have been linked to episodic memory processes in various studies. Both oscillations seem to be vital for processes guided by the medial temporal lobe, such as the retrieval of information from memory. While theta oscillations increase with successful memory, it is unclear what the unique contribution of theta is to various subcomponents of memory. On the other hand, memory-related gamma oscillations have been mainly reported in the hippocampus, leaving the role of neocortical gamma in memory underexplored. In the current study, we explored how unique variability in memory accuracy and memory confidence contributes to fluctuations in theta and gamma power. To this end, we recorded EEG from 54 participants while they performed a source memory task. From this task we obtained their item memory accuracy, source memory accuracy, item memory confidence, and source memory confidence. These behavioral measures were put in a trial-by-trial linear mixed effects model to uncover their unique contribution to the oscillatory power in frontal and parietal regions. Our results are in line with the involvement of theta oscillations in both memory accuracy and confidence, but seem to indicate a main role for theta oscillations in memory-related confidence. In addition, we found that gamma oscillations play various roles in memory-processing, dependent of brain region.

## 1. Introduction

Research into episodic memory, the memory for events that are linked to a specific time and place, frequently looks at two specific subcomponents, referred to as ‘item memory’ and ‘source memory’. Here, item memory reflects memories for specific and unconnected items, whereas source memory represents associative memories that are embedded in a context or encoding source. These two components can be seen as two variants of objective memory accuracy, as item and source memory are often classified as either correct or incorrect. On the other hand, the term ‘memory confidence’ is used to reflect the subjective feeling of confidence we have in a recalled memory. Often this is a graded measure and is correlated with the quality and amount of information retrieved. Memory confidence can be measured in addition to the objective item or source memory performance, to quantify the subjective quality of the memory. When exploring the neural underpinnings of episodic memory in general, research points towards theta and gamma oscillations playing a major role in objective and subjective memory retrieval (Griffiths, Martin-Buro, Staresina, & Hanslmayr, 2021; Herweg, Solomon, & Kahana, 2020; Jensen, Kaiser, & Lachaux, 2007; Lega, Burke, Jacobs, & Kahana, 2016; Nyhus & Curran, 2010; Staudigl & Hanslmayr, 2013). Given the established involvement of these oscillations, it warrants taking a closer look at the specific role they play in memory accuracy and confidence.

Theta (3-7Hz) oscillations may be vital to several memory-related processes carried out in the medial temporal lobe (MTL), the main hub for episodic memory. For example, theta in the MTL appears to be important for creating temporal associations between elements, grouping them in one episodic event in memory (Herweg et al., 2020). Theta in neocortical areas, which can be recorded with electroencephalography (EEG) and magnetoencephalography (MEG), has also been linked to memory. Specifically, theta power is higher when items are remembered (hits), as compared to either forgotten (misses) or correctly identified as new (correct rejections) (Chrastil et al., 2022; Duzel et al., 2003; Duzel, Neufang, & Heinze, 2005; Wynn, Daselaar, Kessels, & Schutter, 2019; Wynn, Kessels, & Schutter, 2020). Similar to the role of theta in the MTL, there is a fairly consistent pattern of cortical theta power increasing when extra associative information is recalled (Addante, Watrous, Yonelinas, Ekstrom, & Ranganath, 2011; Gruber, Tsivilis, Giabbiconi, & Muller, 2008; Guderian & Duzel, 2005; Herweg et al., 2016). In addition, it has been shown that theta power is positively correlated with memory confidence (Wynn et al., 2019; Wynn et al., 2020). When looking at these findings, it appears that theta power is involved in the recollection of items, and that this interacts with the associative information and subjectively perceived confidence associated with that item. However, as these memory processes are correlated, it is unclear what their unique relationship with theta oscillations is.

In addition to theta frequency band, gamma (40-80 Hz) oscillations have been proposed to play an important role in memory retrieval. For instance, there is a close association between hippocampal gamma and neocortical oscillations during both memory encoding and retrieval (Griffiths & Jensen, 2023; Griffiths et al., 2019; Hanslmayr, Staresina, & Bowman, 2016; Pacheco Estefan et al., 2019). The involvement of gamma in memory encoding appears to be mainly through theta-gamma phase amplitude coupling (PAC). Specifically, it has been proposed that each gamma cycle represents a memory specific representation, which are superimposed onto different phases of the theta cycle, forming one cohesive memory event (Griffiths et al., 2019; Heusser, Poeppel, Ezzyat, & Davachi, 2016; Karlsson, Lindenberger, & Sander, 2022; Lisman & Idiart, 1995; Lisman & Jensen, 2013; Ursino, Cesaretti, & Pirazzini, 2023). Furthermore, it has been suggested that increased gamma power during memory retrieval reflects hippocampal pattern completion leading to information reinstatement in the neocortex (Griffiths et al., 2019; Staresina et al., 2016). However, how this affects item memory, source memory, and memory confidence is not fully understood. The literature has suggested that hippocampal gamma is (indirectly) related to all three processes (Merkow, Burke, & Kahana, 2015) or selectively related to source memory (Staresina et al., 2016), that frontoparietal gamma is related to source memory (Burgess & Ali, 2002), and that posterior gamma is related to item memory (Gruber et al., 2008; Osipova et al., 2006). Most studies have investigated hippocampal gamma, and mainly its role in memory encoding. Therefore, it is currently unclear how neocortical gamma oscillations during retrieval are related to specific memory subprocesses.

The aim of the current study is to investigate how variability in memory accuracy (item, source) and memory confidence (high, low) relates to oscillatory power in the theta and gamma band. Specifically, our participants performed a memory task where words were encoded in one of two encoding conditions. During a subsequent memory recognition task, we probed their item memory accuracy, item memory confidence, source memory accuracy, and source memory confidence. Throughout the memory task, we recorded their brain activity via EEG. Linear mixed effect models were used to estimate the unique contribution of each of the behavioral measures to the trial-by-trial theta and gamma power during retrieval. Results from this study will inform us on the relative contribution of various memory subprocesses to the variability in theta and gamma oscillations.

## 2. Methods

### 2.1. Participants

Fifty-four healthy adult right-handed volunteers (32 females, 22 males) with a mean age of 20 (*SD* = 1.55) were included in this study. Data analysed here was part of a larger non-invasive brain stimulation (NIBS) study, which included four sessions. Only the data from the first session is used here, where the participants did not receive any NIBS. Four participants were replaced to maintain our intended sample-size of 54, due to not adhering to task instructions (N=2) and failure to complete all experimental sessions (N=2). All had normal or corrected-to-normal vision, were fluent English speakers, right-handed and free from self-reported neurological or psychiatric conditions. Main exclusion criteria were skin disease, metal in their cranium, epilepsy or a family history of epilepsy, history of other neurological conditions or psychiatric disease, heart disease, use of psychoactive medication or substances, and pregnancy. The study was approved by the institutional review board of Bowdoin College, Brunswick, USA, and carried out in accordance with the standards set by the Declaration of Helsinki.

### 2.2. Stimuli

Stimuli were presented on a personal computer screen with a 21-inch monitor. Stimulus presentation and recording of responses were attained using E-Prime 2.0 software (Psychology Software Tools, Pittsburg, PA). The stimulus material consisted of 400 words per session, varying per participant, randomly chosen from a pool of 1778 words, selected from the MRC Psycholinguistic Database (http://websites.psychology.uwa.edu.au/school/MRCDatabase/uwa_mrc.htm). All words in this database are scored on word frequency, familiarity, and concreteness, which combined leads to an ‘imageability’ rating between 100 and 700 (Coltheart, 1981). We only included nouns and adjectives that had an imageability rating of >300.

### 2.3. Procedure

All participants received written and oral information prior to participation but remained naive regarding the aim of the study. Each volunteer provided written informed consent at the beginning of the first session.

In the intentional encoding phase of the memory task, trials began with the presentation of the response options on the bottom of the screen for 80-120 ms (jittered; see Figure 1). Throughout the trial, these response options remained on the screen. Next, the task cue (Place or Pleasant) was presented in the middle of the screen in yellow font for 500 ms, followed by a blank mask for 200 ms, and the presentation of the to-be-encoded capitalized word for 500 ms. The cue informed the participants on the encoding task in the current trial. When the cue “Place” was presented, participants had to conjure up an image of a scene of a spatial environment that relates to the word that was presented right after. For example, for the word “dirty”, they could imagine a dirty scene or place, such as imagining a scene of a garbage dump or a messy room. When the cue “Pleasant” was presented, participants had to pay attention to the meaning of the word that was presented right after and evaluate how pleasant it is. For example, for the word “dirty”, they could imagine that it is “unpleasant”. Following the word offset, participants had 4 seconds to perform the encoding task while a fixation cross was presented on screen. Thereafter, a question mark replaced the fixation cross, and they had 700 ms to indicate how successful they were at completing the encoding task by responding on a keyboard with their dominant, right hand: “H” = unsuccessful, “U” = partially successful, “I” = successful. In total there were 200 trials; 100 words encoded during the place task and 100 words encoded during the pleasantness task.

**Figure 1.**
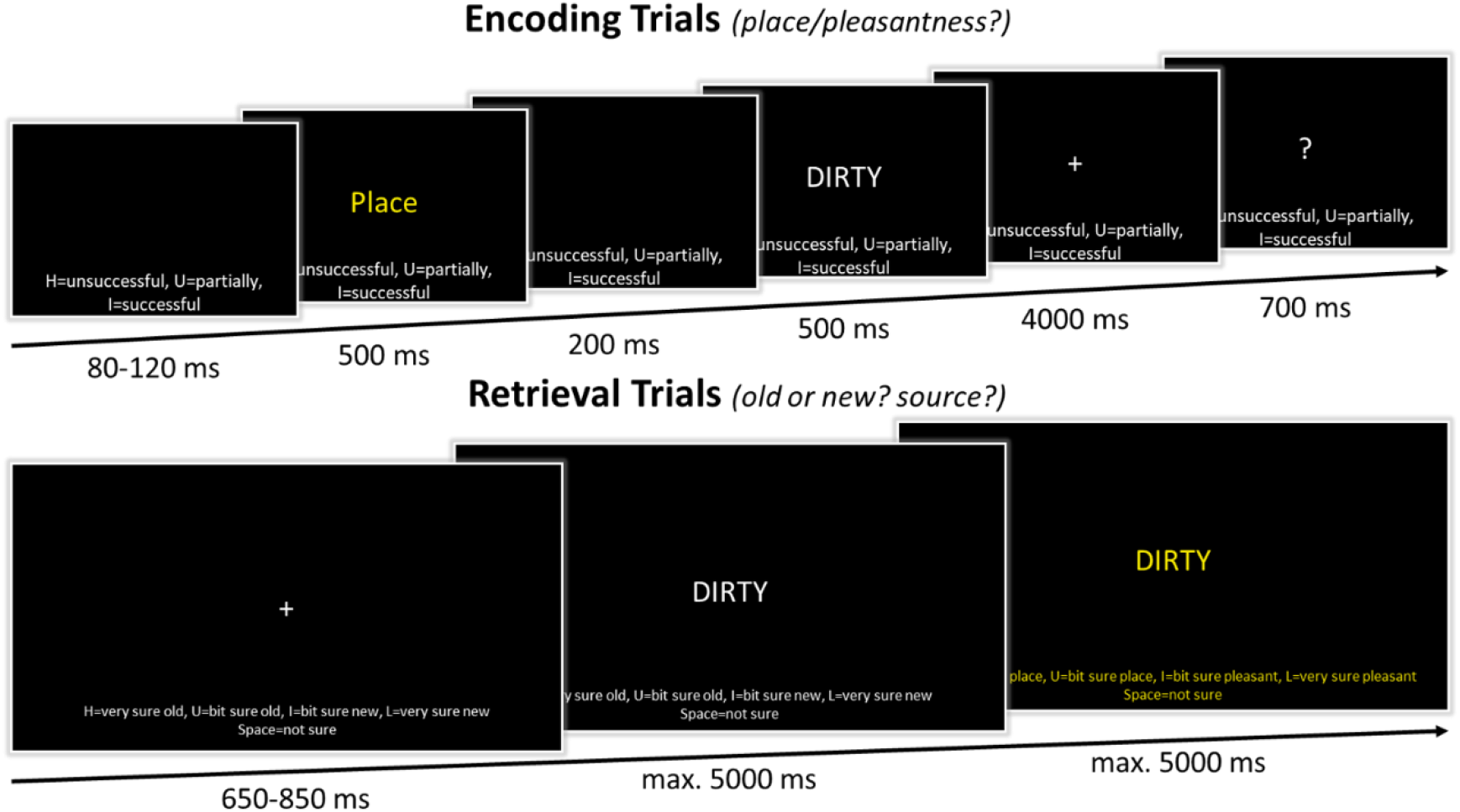
Schematic overview of the memory task. In the encoding phase, participants either had to imagine a spatial scene (place task) or rate the pleasantness (pleasant task) of the presented word. They indicated how successful they were in completing this encoding task. In the retrieval phase, participants first made an old/new response. In the case of an ‘old’ response, participants were asked to indicate what encoding task was performed when first encountering that word.

After the encoding task, participants completed a math task to diminish any rehearsal and recency effect. In this task, mathematical equations (e.g., [56 + (5 + 5) × 2 – 30]) were presented on the computer screen for 15 minutes. Participants were informed that this task was a distraction task, that they should not be nervous about it and just try their best. The math task was self-paced, and participants could alter their answer prior to responding. Responses were attained using the numbers on the keyboard. Results of the math task were not analyzed as it just served as a distraction task.

In the retrieval phase, participants performed a recognition task, including the 200 ‘old’ words that were presented during encoding and 200 ‘new’ words (see Figure 1). Like the encoding trials, the response options were presented on the bottom of the screen throughout the trial. The retrieval trials started with a 650-850 ms (jittered) presentation of a fixation cross. This was followed by a centrally presented capitalized word. Participants were instructed to indicate whether they thought the word was ‘old’ or ‘new’, considering the confidence they had in their decision, on a 5-point scale. Responses were given on a keyboard with their dominant, right hand: “H” = very sure old, “U” = bit sure old, “space bar” = not sure, “I” = bit sure new, “L” = very sure new. Participants had 5 seconds to submit their response and their response would immediately advance the trial. However, to ensure a sufficient time frame for EEG analyses, within the first 800 ms of word presentation, entering a response would not advance the trial immediately. When their response was ‘not sure’, ‘bit sure new’, or ‘very sure new’ the next trial started after their ‘old/new’ decision was finalized. When their response was ‘bit sure old’ or ‘very sure old’, a new screen was presented, and participants could indicate the encoding source of the word. On this screen the source response options were presented: “H” = very sure pleasant, “U” = bit sure pleasant, “space bar” = not sure, “I” = bit sure place, “L” = very sure place. To make it more salient to the participants that a source decision was now required, both the word in the center of the screen and the response options on the bottom were presented in a yellow font. Participants had 5 seconds to indicate their response on the keyboard, after which the next trial began.

To familiarize participants with the memory task, fifteen practice trials preceded both the encoding and retrieval phase of the experiment. Stimuli used during the practice trials were not used in the experimental trials. Following every 100 experimental trials there was a short break of a minimum of 30 seconds. Participants then had 30 seconds to indicate that they were ready to continue.

### 2.4. EEG recording and analyses

EEG was recorded throughout the experimental session. EEG signals were recorded and amplified with an actiCHamp system (Brain Products, Munich, Germany) from sixty-four channels. Amplified analogue voltages (0.1–100 Hz bandpass) were digitized at 10 kHz.

EEG pre-processing and analyses were performed with the use of MATLAB (v2022b, MathWorks Inc., Natrick MA) in combination with the FieldTrip toolbox (Oostenveld, Fries, Maris, & Schoffelen, 2011) and the EEGLAB toolbox (Delorme & Makeig, 2004). Raw signals were downsampled to 1000 Hz and re-referenced to an average reference. The data was high-pass filtered at 0.1 Hz, low-pass filtered at 58 Hz, and an additional band-pass filter was used to remove additional line noise at 60 Hz. Subsequently, the retrieval data was epoched into stimulus-locked time windows. The minimum stimulus presentation duration during retrieval was 800 ms. However, since retrieval was self-paced, stimulus presentation duration varied. For this reason, epochs started 500 ms before stimulus onset and ended either 505 ms before the onset of the following stimulus or after 2000 ms. This resulted in the shortest retrieval epoch being -500 to 1045 ms, and the mean epoch length being -500 to 1810 ms. Epochs with transient muscle or electrode artifacts were rejected based on visual inspection. Additional artifacts were removed using independent component analysis (ICA) in combination with EEGLAB’s ICLabel (Pion-Tonachini, Kreutz-Delgado, & Makeig, 2019). Components classified as muscle artifacts (probability: .9), eye artifacts (probability: .8), heart artifacts (probability: .8), and channel noise artifacts (probability: .9) were removed from the data. A final artifact check was done after ICA by manual inspection.

Spectral power was extracted using Fourier analysis with a 500 ms sliding time window and the application of a Hanning taper. Data was zero-padded and frequencies that were assessed ranged from 1 to 57 Hz in 1 Hz steps. To prepare the data for further statistical analysis, for each trial, spectral data was averaged over the theta band (3-7 Hz) and gamma band (40-50 Hz), a 300 – 800 ms time window, and frontal (F1, F2, F3, F4, F5, F6, Fz) and parietal regions (P1, P2, P3, P4, P5, P6, Pz), based on previous EEG studies (Babiloni et al., 2004; Burgess & Ali, 2002; Gruber et al., 2008; Wynn et al., 2019; Wynn et al., 2020). This gave us four variables: ‘Frontal Theta’, ‘Parietal Theta’, ’Frontal Gamma’, and ‘Parietal Gamma’, which were used in further trial-by-trial analyses.

### 2.5. Statistical analyses

All statistical analyses were performed in RStudio (RStudio version 2023.06.2, R version 4.2.0; R Core Team, 2021). For the behavioral analysis, hit rate, false alarm rate, d-prime and high-confidence rate were calculated as follows:

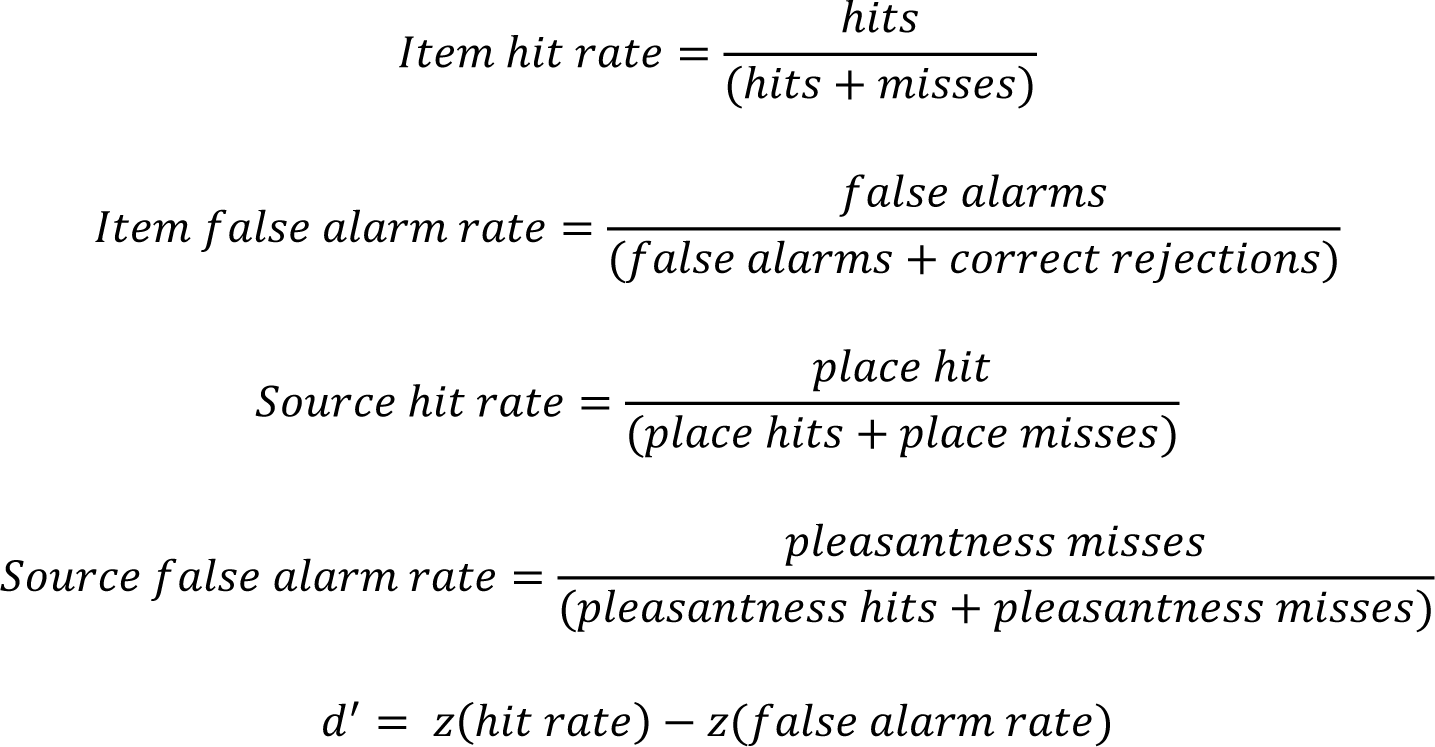

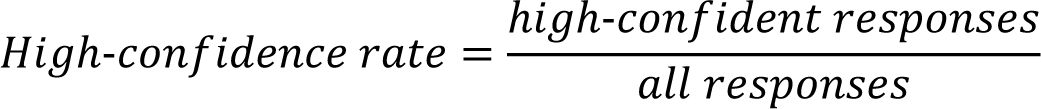

For these analyses, two-sided pairwise t-tests were performed with an alpha of .05. Cohen’s d was used as an effect size of the t-tests.

For all trial-based oscillatory analyses, each trial was coded based on memory status (‘old’ or ‘new’) and memory accuracy (‘correct’ or ‘incorrect’), the combination of these two would make up the following memory categories: hits (‘old’ and ‘correct’), misses (‘old’ and ‘incorrect’), correct rejections (‘new’ and ‘correct’), and false alarms (‘new’ and ‘incorrect’). Due to the low number of ‘a bit sure’ and ‘not sure’ responses, these two responses were combined into one ‘low confidence’ level. Therefore, trial-based memory confidence on every trial was re-coded into ‘low’ and ‘high’ confidence. Outlier-correction was performed on the trial-by-trial EEG data, with a cut-off of three standard deviations above or below the mean. For these analyses, linear mixed effects models were used to predict spectral power in the gamma and theta frequency band from the behavioral responses from each individual trial (lme4 package (v. 1.1.29); Bates, Mächler, Bolker, & Walker, 2015). A mixed effect model was deemed most appropriate as it can account for within- and between-subject variability, through employing by-participant varying intercepts and slopes. Specifically, the following models were used in the analyses:

Item memory:

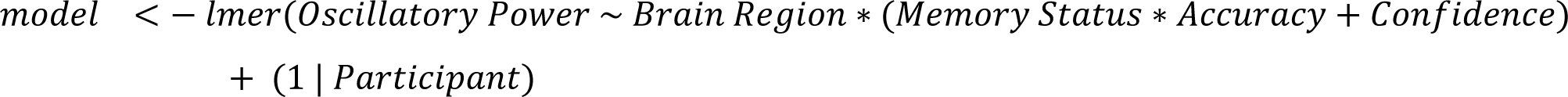

Source memory

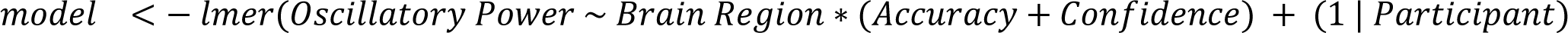

The fixed effects were Brain Region (frontal, parietal), Memory Status (old, new), Accuracy (correct, incorrect), and Confidence (high, low). As we were interested in two specific brain regions, we let Brain Region interact with the behavioral responses. Given that in item memory, we were interested in specific combinations of Memory Status and Accuracy, we also let those predictors interact. The combinations of interest in the current study were old and correct (hits), new and correct (correct rejections), old and incorrect (misses), and new and incorrect (false alarms). In addition, we had a by-participant varying intercept and slope, to consider individual differences. All categorical fixed effects were sum-coded, and the linear outcome measure was standardized. These models were run separately for theta and gamma power, leading to four models used in total. Significance of the model outputs were generated by the lmerTest package (v. 3.1.3; Kuznetsova, Brockhoff, & Christensen, 2017), which applies the Satterthwaite method for estimating degrees of freedom, with an alpha level of .05. In the case of a significant interaction, pairwise comparisons on the estimated marginal means were used to further investigate this. These comparisons were restricted to the following comparisons: hits versus correct rejections, hits versus misses, correct rejections versus false alarms, and high-confidence versus low-confidence. The first three were used to inform us on the effects of accuracy, while controlling for confidence, while the last one was used to look at the effect of confidence, while controlling for accuracy.

## 3. Results

### 3.1. Behavioral results

During encoding, participants successfully thought of either a place or the pleasantness regarding the presented word. The average imaging success was 87.37% (*SD* = 8.40), with no significant difference between the Pleasantness (*M* = 87.35%, *SD* = 10.07) and Place (*M* = 87.15%, *SD* = 8.96) conditions (*t*(53) = 0.18, *p* = .86, *d* = .022). After the fixed four second time window, their average imaging reaction time (RT) was 378 ms (*SD* = 47), with a significantly faster response in the Pleasantness (*M* = 373, *SD* = 47) than Place (*M* = 383, *SD* = 48) condition (*t*(53) = -3.75, *p* < .001, *d* = -.22).

For the full description of the retrieval behavioral measures, see Table 1. The average item memory performance, as quantified by d-prime, was 2.31 (*SD* = 0.59) and the average RT, in ms, for the old/new judgement was 1642 (*SD* = 309). RTs showed a significant difference between the correct and incorrect old/new responses (*t*(53) = -12.06, *p* < .001, *d* = -.94). The average source memory performance, as quantified by d-prime, was 1.64 (*SD* = 0.73) and the average RT for the source judgement was 1273 (*SD* = 447). RTs showed a significant difference between correct and incorrect source responses (*t*(53) = -3.09, *p* = .003, *d* = -.33). The results indicated that overall, the Pleasantness and Place conditions were not significantly different, therefore they were combined in further analyses (see Table 1).

**Table 1.** Mean values of behavioral performance during memory retrieval, with the standard deviation in brackets. Significant differences between Place and Pleasantness conditions are indicated in the table.

|  | Item Memory |  |  | Source memory |  |  |
| --- | --- | --- | --- | --- | --- | --- |
|  | All | Place | Pleasantness | All | Place | Pleasantness |
| Hit Rate | .77 (.093) | .76 (.097) | .78 (.10) | .72 (.15) | - | - |
| False Alarm Rate | .08 (.065) | - | - | .18 (.11) | - | - |
| D-prime | 2.31 (0.59) | 2.29 (0.59) | 2.34 (0.63) | 1.64 (.73) | - | - |
| HC Responses (%) | .61 (.21) | - | - | .54 (.22) | - | - |
| HC Hits (%) | .83 (.13) | .83 (.13) | .83 (.14) | .68 (.22) | .72 (.22) | .63 (.24) * |
| HC Correct Rejections (%) | .52 (.30) | - | - | - (-) |  |  |
| RT All Trials (in ms) | 1642 (309) | 1663 (333) | 1664 (320) | 1273 (447) | 1256 (452) | 1284 (472) |
| RT Correct Trials (in ms) | 1587 (292) | 1646 (342) | 1626 (323) | 1258 (503) | 1242 (516) | 1292 (535) |
| RT Incorrect Trials (in ms) | 1911 (392) | 1779 (431) | 1845 (424) | 1423 (504) | 1451 (559) | 1412 (506) |

HC = High-Confident, RT = Reaction Time, \* = p-value < .05

### 3.2. Oscillatory results

When investigating how elements of item memory predict theta power, the model showed that there was a significant Brain Region x Memory Status x Accuracy interaction (see Figure 2 and Table 2). Post-hoc tests (see Table 3) on the estimated marginal means revealed that predicted theta power was higher for hits, as compared to correct rejections and misses, in the frontal and parietal regions. In addition, parietal theta power was significantly higher in high-confident responses, as compared to low-confident responses. This indicates that theta oscillations play a role in both item memory accuracy and item memory confidence, where parietal theta may be more tuned to confidence judgements, as compared to frontal theta.

**Figure 2.**
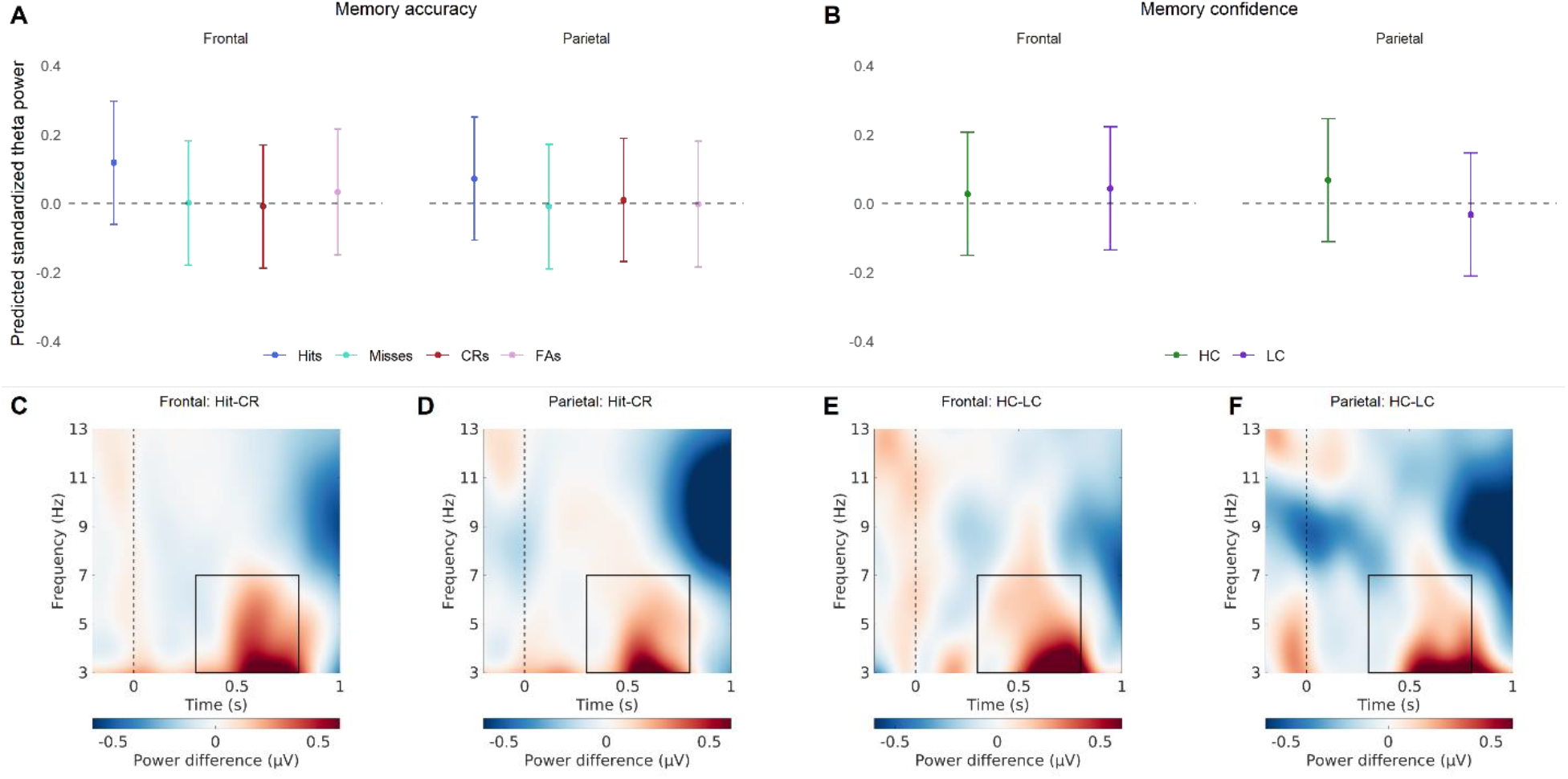
The relationship between theta power and item memory. (A) Predicted standardized power obtained from the linear mixed effect model for the memory accuracy conditions (hits, misses, correct rejections, false alarms), for frontal and parietal regions. The values shown reflect predicted theta while controlling for the other predictors in the model, like memory confidence (B) Predicted standardized power obtained from the linear mixed effect model for the memory confidence conditions (high-confidence, low-confidence), for frontal and parietal regions. The values shown reflect predicted theta while controlling for the other predictors in the model, like memory accuracy. (C) The difference in power between hits and correct rejections for the frontal EEG channels. (C) The difference in power between hits and correct rejections for the parietal EEG channels. (C) The difference in power between high- and low-confidence for the frontal EEG channels. (D) The difference in power between high- and low-confidence for the Parietal EEG channels. The frequency and time window used for the models is indicated in the lower plots by a rectangle.

**Table 2.** Theta - Item Memory: Model.

|  | Estimate | Std. Error | DF | T-value | P-value |
| --- | --- | --- | --- | --- | --- |
| Intercept | -0.006 | 0.092 | 56 | -0.063 | 0.950 |
| Brain Region | 0.046 | 0.015 | 36531 | 3.162 | 0.002* |
| Memory Status | -0.020 | 0.019 | 36536 | -1.040 | 0.299 |
| Accuracy | -0.016 | 0.017 | 36539 | -0.939 | 0.348 |
| Confidence | 0.042 | 0.010 | 36560 | 4.225 | 0.000* |
| Memory Status * Accuracy | 0.114 | 0.021 | 36538 | 5.423 | 0.000* |
| Brain Region * Memory Status | -0.012 | 0.019 | 36531 | -0.655 | 0.512 |
| Brain Region * Accuracy | -0.027 | 0.016 | 36531 | -1.688 | 0.091<sup>+</sup> |
| Brain Region * Confidence | -0.058 | 0.009 | 36531 | -6.483 | 0.000 |
| Brain Region * Memory Status * Accuracy | 0.045 | 0.021 | 36531 | 2.171 | 0.030* |

**Table 3.**
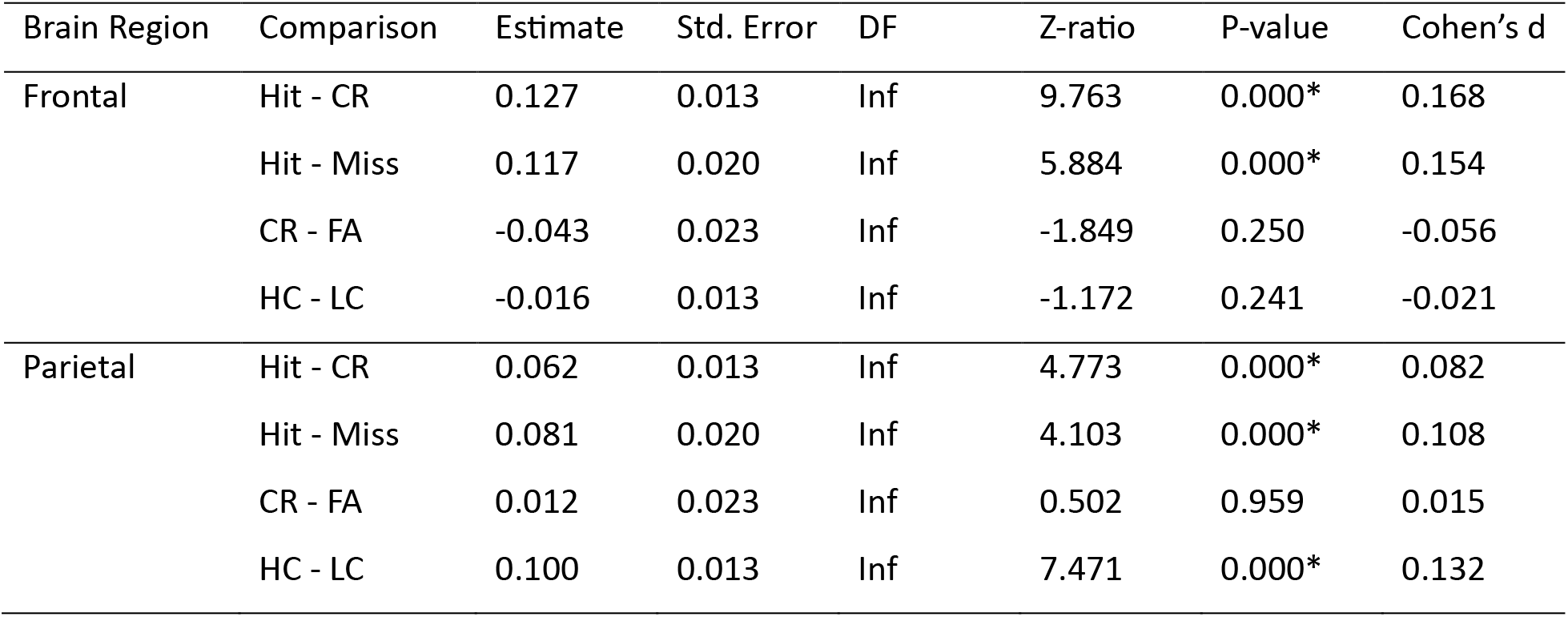
Theta - Item Memory: Pairwise Comparisons.

Regarding the relationship between theta power and source memory, the model showed that there was a marginal Brain Region x Accuracy interaction, and a significant main effect of Confidence (see Figure 3 and Table 4). Predicted theta was higher for high-confident source decisions (*M* = 0.13, *SE* = 0.095), as compared to low-confident source decisions (*M* = 0.062, *SE* = 0.095). This indicates that regarding source memory, there is evidence for a role of theta in confidence in both parietal and frontal regions.

**Figure 3.**
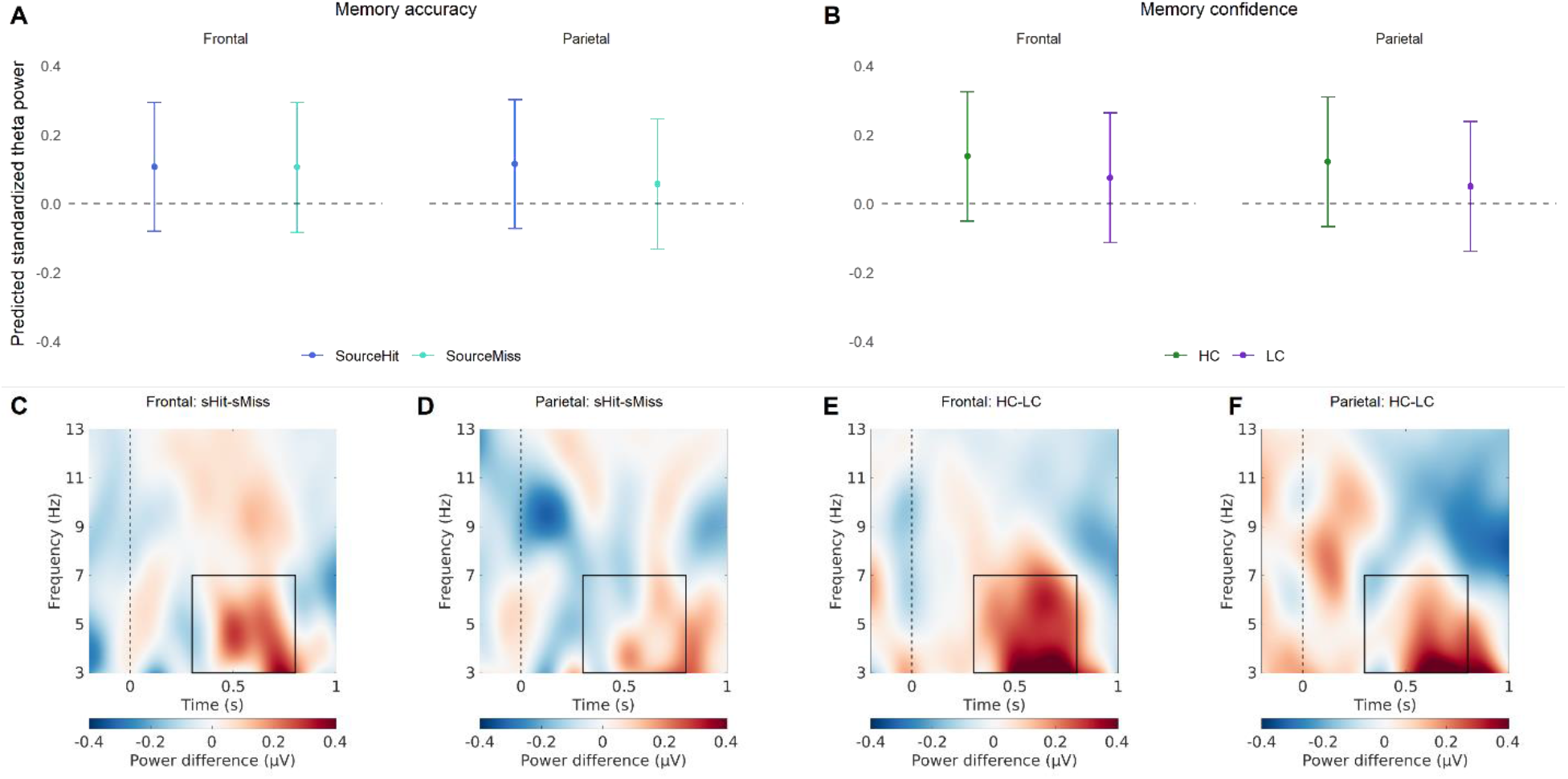
The relationship between theta power and source memory. (A) Predicted standardized power obtained from the linear mixed effect model for the memory accuracy conditions (source hits, source misses), for frontal and parietal regions. The values shown reflect predicted theta while controlling for the other predictors in the model, like memory confidence (B) Predicted standardized power obtained from the linear mixed effect model for the memory confidence conditions (high-confidence, low-confidence), for frontal and parietal regions. The values shown reflect predicted theta while controlling for the other predictors in the model, like memory accuracy. (C) The difference in power between source hits and source misses for the frontal EEG channels. (C) The difference in power between source hits and source misses for the parietal EEG channels. (C) The difference in power between high- and low-confidence for the frontal EEG channels. (D) The difference in power between high- and low-confidence for the Parietal EEG channels. The frequency and time window used for the models is indicated in the lower plots by a rectangle.

**Table 4.** Theta - Source Memory: Model.

|  | Estimate | Std. Error | DF | T-value | P-value |
| --- | --- | --- | --- | --- | --- |
| Intercept | 0.048 | 0.096 | 55 | 0.497 | 0.621 |
| Brain Region | 0.026 | 0.013 | 14021 | 1.970 | 0.049* |
| Accuracy | 0.030 | 0.016 | 14031 | 1.798 | 0.072<sup>+</sup> |
| Confidence | 0.066 | 0.016 | 14048 | 4.157 | 0.000* |
| Brain Region * Accuracy | -0.027 | 0.016 | 14021 | -1.729 | 0.084<sup>+</sup> |
| Brain Region * Confidence | -0.005 | 0.014 | 14021 | -0.328 | 0.743 |

The model looking at the relationship between item memory and gamma power showed that there was a significant Brain Region x Accuracy interaction, and a significant Brain Region x Confidence interaction (see Figure 4 and Table 5). Post-hoc tests (see **Error! Reference source not found.**) on the estimated marginal means revealed that the predicted parietal gamma power is significantly higher for hits, as compared to both correct rejections and misses. In addition, predicted frontal gamma power was significantly higher in Low-Confident responses, as compared to High-confident responses. This indicates that gamma oscillations play a role in both item memory accuracy and item memory confidence, but that this is specific to parietal or frontal regions, respectively.

**Figure 4.**
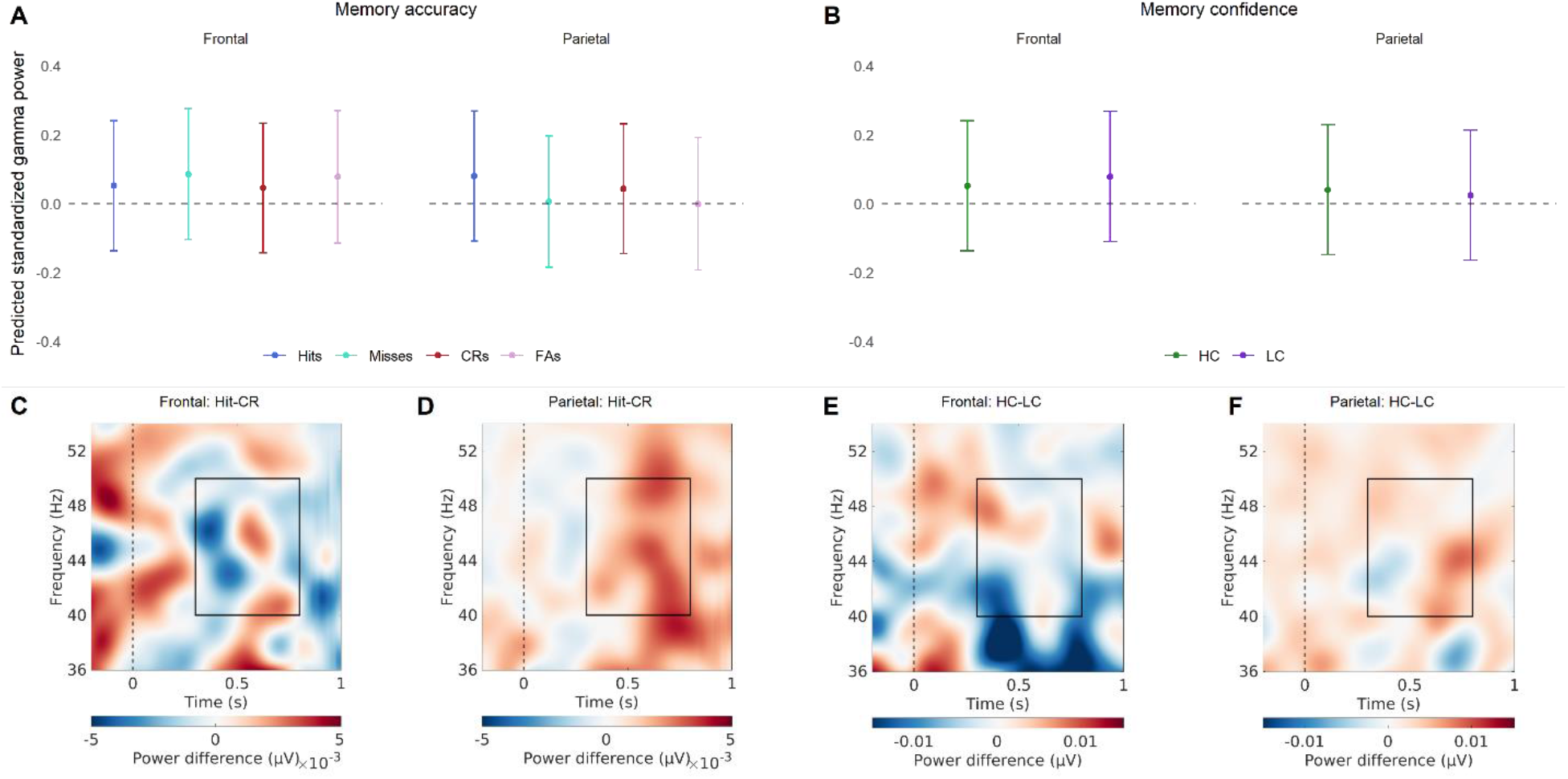
The relationship between gamma power and item memory. (A) Predicted standardized power obtained from the linear mixed effect model for the memory accuracy conditions (hits, misses, correct rejections, false alarms), for frontal and parietal regions. The values shown reflect predicted gamma while controlling for the other predictors in the model, like memory confidence (B) Predicted standardized power obtained from the linear mixed effect model for the memory confidence conditions (high-confidence, low-confidence), for frontal and parietal regions. The values shown reflect predicted gamma while controlling for the other predictors in the model, like memory accuracy. (C) The difference in power between hits and correct rejections for the frontal EEG channels. (C) The difference in power between hits and correct rejections for the parietal EEG channels. (C) The difference in power between high- and low-confidence for the frontal EEG channels. (D) The difference in power between high- and low-confidence for the Parietal EEG channels. The frequency and time window used for the models is indicated in the lower plots by a rectangle.

**Table 5.** Gamma - Item Memory: Model.

|  | Estimate | Std. Error | DF | T-value | P-value |
| --- | --- | --- | --- | --- | --- |
| Intercept | 0.041 | 0.097 | 55 | 0.425 | 0.672 |
| Brain Region | 0.050 | 0.014 | 36443 | 3.460 | 0.001* |
| Memory Status | 0.007 | 0.019 | 36448 | 0.376 | 0.707 |
| Accuracy | 0.006 | 0.016 | 36450 | 0.353 | 0.724 |
| Confidence | -0.006 | 0.010 | 36469 | -0.590 | 0.555 |
| Memory Status * Accuracy | 0.014 | 0.021 | 36449 | 0.690 | 0.490 |
| Brain Region * Memory Status | 0.001 | 0.018 | 36443 | 0.048 | 0.962 |
| Brain Region * Accuracy | -0.038 | 0.016 | 36443 | -2.429 | 0.015* |
| Brain Region * Confidence | -0.022 | 0.009 | 36443 | -2.471 | 0.013* |
| Brain Region * Memory Status * Accuracy | -0.016 | 0.020 | 36443 | -0.774 | 0.439 |

**Table 6.**
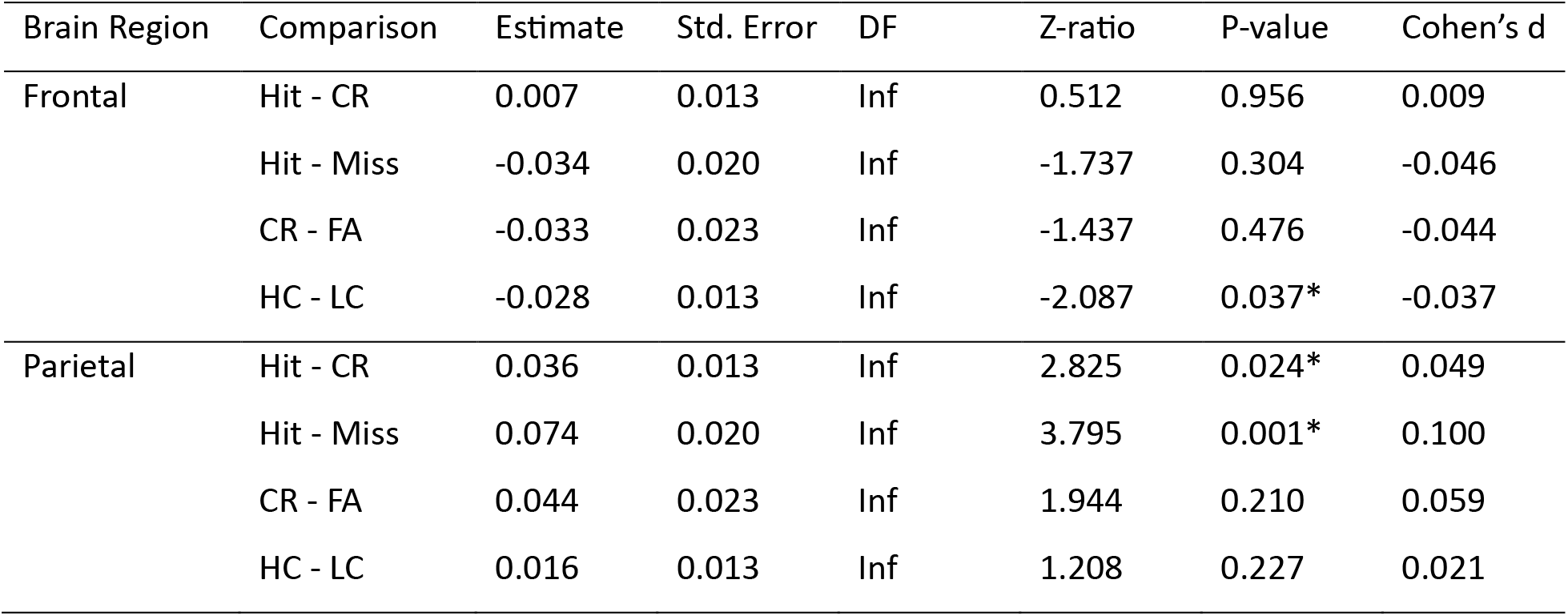
Gamma - Item Memory: Pairwise Comparisons.

Regarding the relationship between gamma power and source memory, the model showed that there was a significant Brain Region x Confidence interaction and a significant main effect of Accuracy (see Figure 5 and Table 7). Predicted gamma was lower for correct source decisions (*M* = 0.084, *SE* = 0.097) than incorrect source decisions (*M* = 0.051, *SE* = 0.097). Post-hoc tests (see Table 8) on the estimated marginal means revealed that in the frontal region, predicted gamma power was marginally higher for low-confident responses, as compared to high-confident responses. The opposite pattern was found in the parietal region, there predicted gamma power was significantly higher for high-confident responses, as compared to low-confident responses. This indicates that gamma plays a role in both source memory accuracy and confidence.

**Figure 5.**
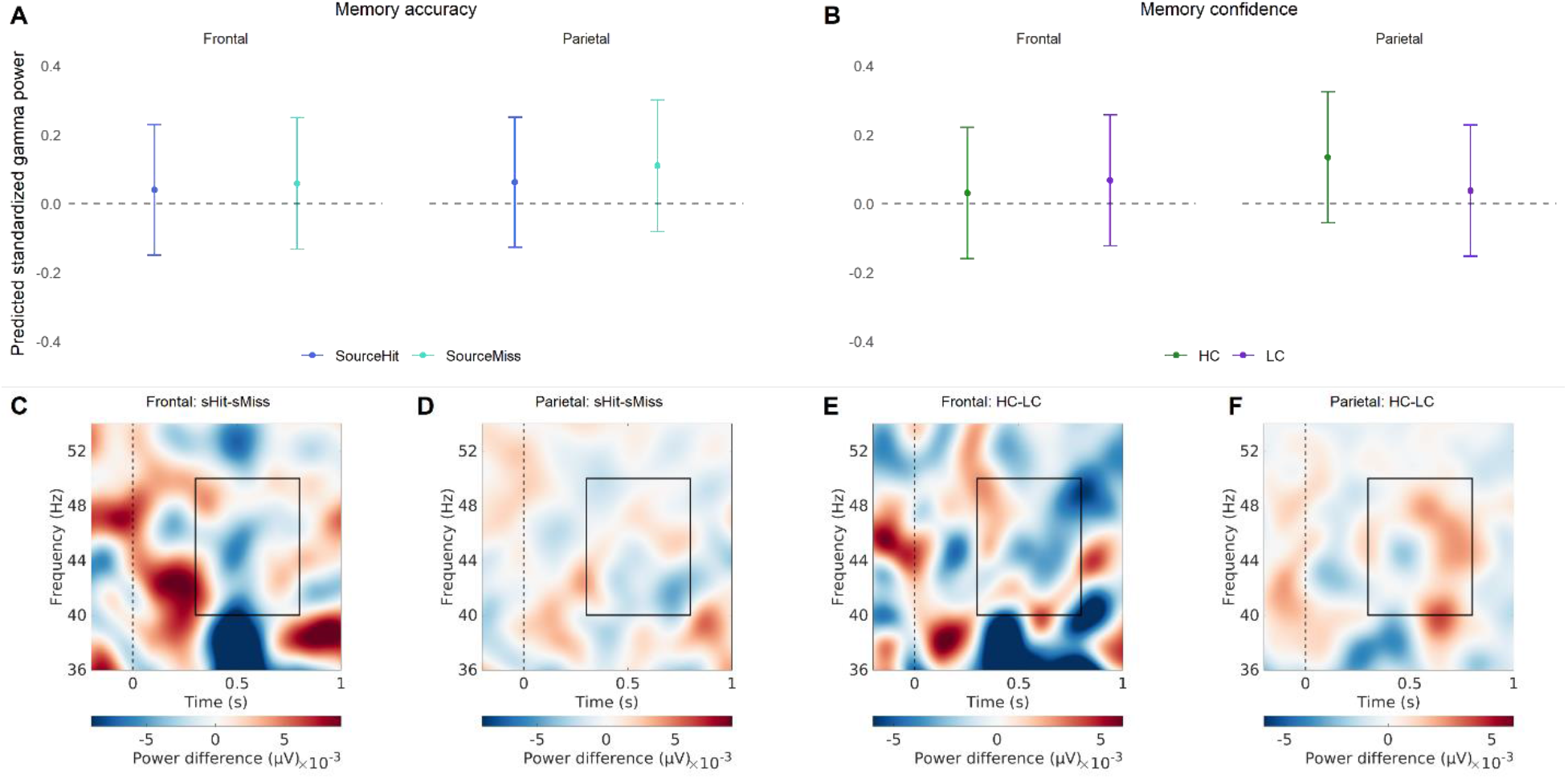
The relationship between gamma power and source memory. (A) Predicted standardized power obtained from the linear mixed effect model for the memory accuracy conditions (source hits, source misses), for frontal and parietal regions. The values shown reflect predicted gamma while controlling for the other predictors in the model, like memory confidence (B) Predicted standardized power obtained from the linear mixed effect model for the memory confidence conditions (high-confidence, low-confidence), for frontal and parietal regions. The values shown reflect predicted gamma while controlling for the other predictors in the model, like memory accuracy. (C) The difference in power between source hits and source misses for the frontal EEG channels. (C) The difference in power between source hits and source misses for the parietal EEG channels. (C) The difference in power between high- and low-confidence for the frontal EEG channels. (D) The difference in power between high- and low-confidence for the Parietal EEG channels. The frequency and time window used for the models is indicated in the lower plots by a rectangle.

**Table 7.** Gamma - Source Memory: Model.

|  | Estimate | Std. Error | DF | T-value | P-value |
| --- | --- | --- | --- | --- | --- |
| Intercept | 0.069 | 0.097 | 55 | 0.711 | 0.480 |
| Brain Region | 0.008 | 0.013 | 14013 | 0.609 | 0.543 |
| Accuracy | -0.033 | 0.016 | 14021 | -2.112 | 0.035* |
| Confidence | 0.030 | 0.015 | 14038 | 1.957 | 0.050<sup>+</sup> |
| Brain Region * Accuracy | 0.015 | 0.015 | 14013 | 0.958 | 0.338 |
| Brain Region * Confidence | -0.067 | 0.014 | 14013 | -4.842 | 0.000* |

**Table 8.** . Gamma - Source Memory: Pairwise Comparisons.

| Brain Region | Comparison | Estimate | Std. Error | DF | Z-ratio | P-value | Cohen's d |
| --- | --- | --- | --- | --- | --- | --- | --- |
| Frontal | HC - LC | -0.037 | 0.021 | Inf | -1.793 | 0.073<sup>+</sup> | -0.049 |
| Parietal | HC - LC | 0.097 | 0.021 | Inf | 4.697 | 0.000* | 0.130 |

## Discussion

This EEG study explored how variability in objective and subjective item and source memory influences theta and gamma oscillations. By utilizing a source memory task that incorporates confidence ratings, we were able to investigate item and source memory and their respective accuracy and confidence levels. A model-based approach was used to disentangle the unique contributions of each of these behavioral measures to the trial-by-trial theta and gamma power during retrieval.

When we focus on theta oscillations, our results show that most of the behavioral measures we investigated influenced theta power measured during retrieval (see Table 9), which is in line with previous literature (Addante et al., 2011; Duzel et al., 2005; Gruber et al., 2008; Wynn et al., 2019; Wynn et al., 2020). These studies showed greater theta power for hits than correct rejection and misses, and greater theta power for high-confident than low-confident responses. Here, we replicate these findings and show that these findings remain unchanged when other correlated variables are kept constant. Specifically, when controlling for memory confidence, we found memory accuracy effects, and when controlling for memory accuracy, we found memory confidence effects. The association between item confidence and theta was found to be specific to the parietal region. This is consistent with previous studies which have linked parietal theta to memory confidence, irrespective of memory status (Wynn et al., 2019; Wynn et al., 2020). However, it is of note that we did not find a relationship between theta power and source memory accuracy, given the literature showing this relation (Addante et al., 2011; Gruber et al., 2008; Guderian & Duzel, 2005; Herweg et al., 2016). In this study the role of theta in source memory seemed to be specific to source memory confidence, a measure which is not often included in analyses. Our findings thus suggest that previous associations between theta power and source memory accuracy, may have been mediated by a confidence effect.

**Table 9.**
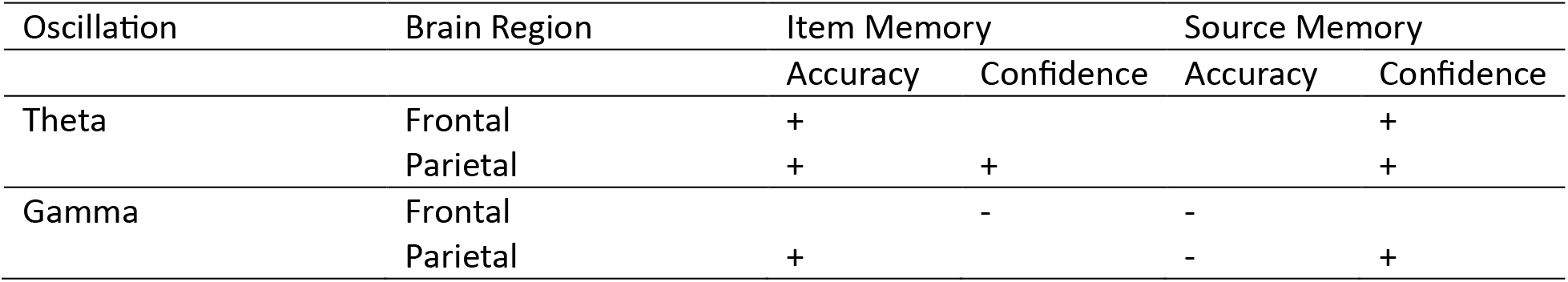
Summary of relationships between oscillatory power and behavioral measurements. Significant positive relationships indicated with an “+” and significant negative ones with an “-”.

The theta effects we found on memory accuracy seemed to be specific to retrieving information from memory, as compared to novelty detection. Given that theta power differentiated between correct and incorrect responses for old items, but not new items. This seems to contradict previous literature reporting a link between evoked frontal theta and novelty processing (Wynn et al., 2019; Wynn et al., 2020). However, these studies showed an interaction between the novelty effect and memory confidence, not a main effect. Here we did not investigate any interactions between memory accuracy and confidence since our main focus was disentangling the unique effects of behavioral measures. Furthermore, previous findings were specific to evoked theta, while here no distinction was made between evoked and induced theta. This could indicate that novelty effects may be specific to time- and phase-locked theta oscillations. Another explanation for these findings is that the main role of theta power pertains to memory confidence, which mediates subsequent novelty decisions in a similar way it can mediate source memory decisions. Overall, our theta results are in concordance with previous literature showing a link between theta power, and both memory accuracy and confidence. This study additionally provides evidence for independent relations between these memory measures and retrieval-related theta oscillations. Moreover, it supports the hypothesis that theta power plays a significant role in memory-related decision-making, given the positive relationship with memory confidence.

Our findings also add evidence for a link between retrieval-related processes and neocortical gamma oscillations (see Table 9). Previous studies have linked gamma oscillations to item and source memory accuracy (Burgess & Ali, 2002; Gruber et al., 2008) and decision-making processes (Castelhano, Duarte, Wibral, Rodriguez, & Castelo-Branco, 2014; Polanía, Krajbich, Grueschow, & Ruff, 2014). Our results are in accordance with this by showing a link between parietal gamma and both item memory accuracy and source memory confidence. The latter supports the involvement in decision-making processes regarding source memories. Interestingly, the link between gamma oscillations and memory confidence seems to differ between frontal and parietal regions. In the frontal regions, there was a negative relationship, while there was a positive one in the parietal regions. When looking at Figure 4C/E and Figure 5C/E we can see that in the frontal gamma difference plot includes negative and positive values, dependent on the specific gamma frequency and time window. This might indicate multiple processes underlying frontal gamma, each having different effects on source memory. It has been proposed before that gamma oscillations do not reflect a unitary cognitive process and sub-bands serve different functions (Bi, Segneri, Di Volo, & Torcini, 2020; Castelhano et al., 2014; Fernández-Ruiz et al., 2021; Vivekananda et al., 2021). Additional research is needed to shed more light on this and to confirm that the negative effect found here is consistently the most dominant one. The positive relation between gamma and source confidence in the parietal region seems to fit with the patterns found in previous literature (Burgess & Ali, 2002; Castelhano et al., 2014; Polanía et al., 2014). In addition, consistent with our results regarding theta power, we found no evidence for a link between gamma oscillations and novelty processing. Overall, our results add to the sparse literature on the role of neocortical gamma and memory processing, by showing a link between gamma power and all the memory-related behavioral measures.

To summarize, we utilized a trial-by-trial model-based approach to uncover the unique relationships between behavioral memory measures and neuronal oscillations. Our results indicate that theta and gamma oscillations are linked to both memory accuracy and confidence. However, whereas theta oscillations seem to primarily play a role in memory-related confidence, gamma oscillations appear to have multiple functions in memory-processing, dependent on brain area.

## Acknowledgements

This work was supported by the National Institutes of Health (NIH) grant R15MH114190.

